# An Open-Source Magnetofluorescence Imaging Platform for Plate-Scale Screening of Magnetic Field Effects in Live Bacteria

**DOI:** 10.64898/2026.07.28.741305

**Authors:** Alessandro Lodesani, Brian L. Ross, Venkatesh Sridharan, Clarice D. Aiello

## Abstract

Magnetic field effects (MFEs) in biological systems are typically small and experimentally challenging to measure reproducibly across large sample populations. Existing approaches to measure such effects often rely on low-throughput microscopy or custom-built magnetic stimulation systems that provide limited control over magnetic field geometry, synchronization, or experimental automation. Here, we present an open-source magnetofluorescence imaging platform designed for bacterial plate-scale screening of MFEs in live colonies. The instrument integrates a programmable three-axis vector electromagnet, synchronized fluorescence excitation and imaging, and integrated acquisition software with per-frame metadata logging on a hardware-synchronized data acquisition card. An extensive calibration procedure enables accurate generation of arbitrary magnetic field vectors, while synchronized triggering ensures deterministic alignment between field application, illumination, and image acquisition. The system images an entire 100 mm Petri dish in a single acquisition. Typical experiments monitor hundreds of bacterial colonies simultaneously over multi-hour acquisition sequences. Control software, calibration routines, mechanical design files, and acquisition workflows are provided openly to facilitate replication. Instrument performance is demonstrated through detection of magnetic field-dependent fluorescence changes in E. coli expressing the engineered magnetosensitive fluorescent protein MagLOV2. This instrument provides a flexible and scalable platform for high-throughput magnetobiology, synthetic biology, and quantum biology experiments.

## 1. Introduction

Magnetic field effects (MFEs) in biological systems have attracted increasing attention across quantum biology, biophysics, and synthetic biology due to accumulating evidence that weak magnetic fields can influence a myriad of biochemical and cellular processes under certain experimental conditions [1–4]. Proposed mechanisms of biological magnetosensing include radical-pair spin dynamics, spin-dependent reaction pathways, and magnetically sensitive photochemistry in flavin-containing proteins [5, 6]. According to the radical pair mechanism theory [7, 8], fluorescence intensity depends on the spin of electrons in a radical pair and can be modulated by changing the magnetic field environment, making this quantity a proxy for measuring spin states. Experimental studies have reported MFEs in systems ranging from avian magnetoreception and cryptochrome photochemistry [9–11] to reactive oxygen species production [12], cellular metabolism [4], and engineered magnetosensitive proteins [13–16].

Despite growing interest in the field, experimental investigation of MFEs remains technically challenging. Reported effects are often small in magnitude, sensitive to illumination conditions and magnetic field geometry, and difficult to reproduce across laboratories. A major practical limitation is the lack of standardized instrumentation capable of integrating precise magnetic field control with high-throughput optical readout. Many existing experiments rely on modified microscopy systems, hand-positioned permanent magnets, or custom-built coil assemblies with limited automation and limited control over magnetic field direction and temporal modulation. These constraints make systematic screening of genetically diverse samples difficult and reduce experimental reproducibility.

Recent advances in synthetic biology and protein engineering have created a growing need for scalable platforms capable of screening magnetosensitive phenotypes across large numbers of samples. Colony-scale fluorescence imaging is particularly attractive because it enables simultaneous monitoring of hundreds of spatially separated bacterial colonies under closely matched environmental conditions. Compared to single-cell microscopy approaches, this strategy sacrifices spatial resolution in favor of substantially increased throughput and statistical power, making it well-suited for screening applications in magnetobiology.

Here we present an open-source magnetofluorescence imaging platform designed specifically for fluorescence image-based scalable screening of MFEs in living colonies. The instrument combines a large-field fluorescence imaging system with a programmable three-axis vector electromagnet implemented through hardware-synchronized control. The platform supports arbitrary static and dynamic magnetic field waveforms, integrated acquisition software with per-frame metadata logging, and reproducible calibration workflows. The system was designed with accessibility and extensibility in mind: all control software, calibration tools, mechanical design files, and experiment workflows are openly available.

To demonstrate instrument performance, we report measurements of magnetic field-dependent fluorescence changes in *E. coli* expressing the engineered magnetosensitive fluorescent protein MagLOV2 [15]. More broadly, the instrument is intended to serve as a flexible platform for magnetosensitivity screening experiments and for systematic exploration of magnetic field interactions in biological and bioengineered systems.

## 2. Instrument Overview

Figure 1 provides a system-level overview of the architecture and signal flow. The instrument integrates fluorescence imaging, programmable magnetic field generation, and synchronized hardware control into a single automated platform. Python-based control software running on a host computer coordinates image acquisition and magnetic field control through a data acquisition card. The data acquisition card generates synchronized trigger and control signals for the camera, LED driver, and three-axis electromagnet controller, while also acquiring analog readout from a Lakeshore F71 magnetometer during calibration and validation procedures. Fluorescence excitation is delivered from above the sample using a high-power LED source, while emission is collected in an epifluorescence geometry by a scientific sCMOS camera positioned directly above the sample stage. The sample itself is positioned above the center of a water-cooled vector electromagnet capable of generating arbitrary magnetic field vectors under software control. A recirculating chiller provides thermal management for sustained electromagnet operation. This modular architecture enables precise synchronization between magnetic field waveforms, illumination timing, and image acquisition throughout an experiment.

**Fig. 1.**
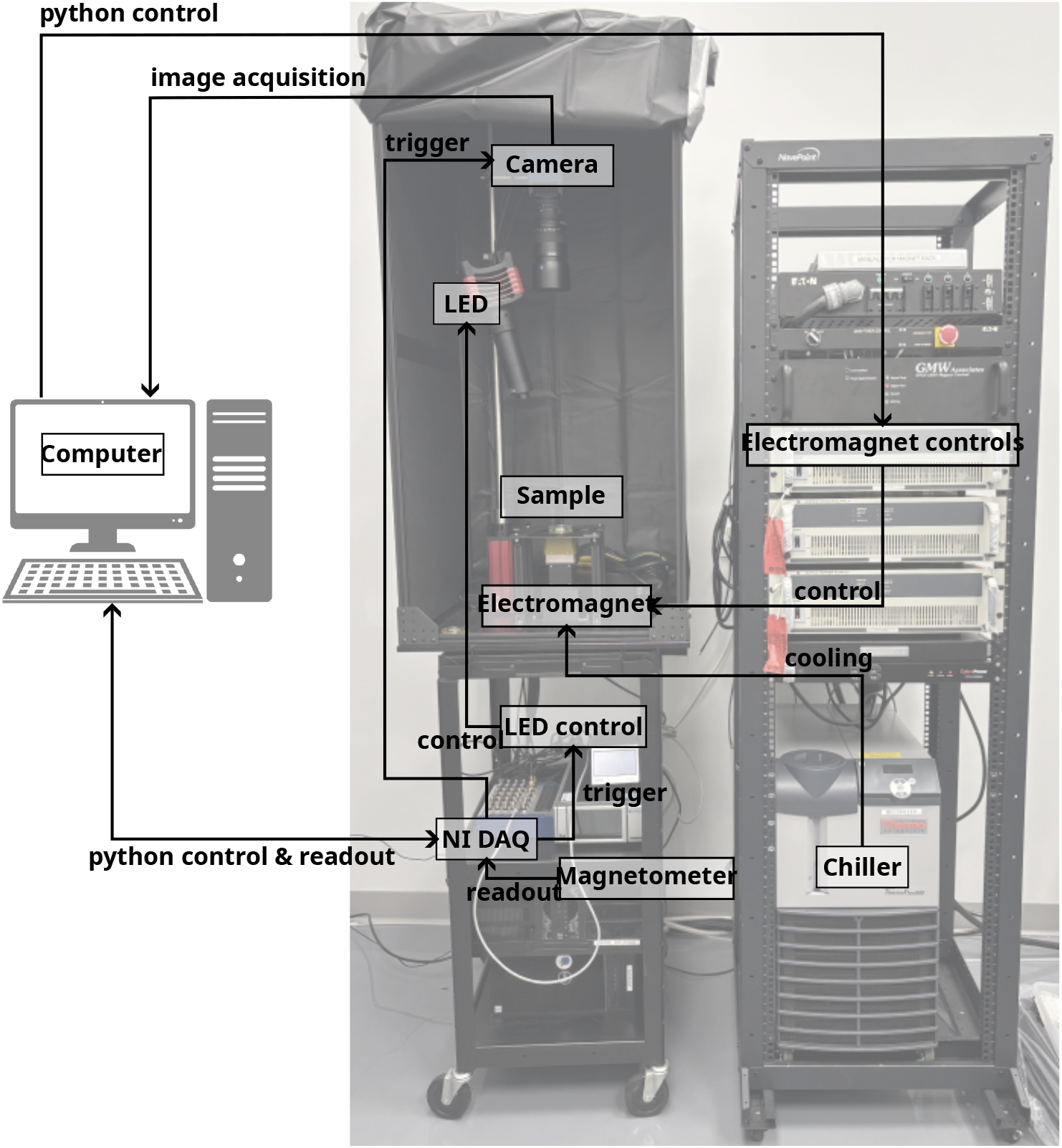
Instrument schematic. The system integrates an epifluorescence imaging module with a programmable vector electromagnet, synchronized LED illumination, and computer-controlled data acquisition. A Python-based control software coordinates magnetic field generation, LED triggering, camera acquisition, and magnetometer readout through a National Instruments DAQ, enabling fully automated magnetofluorescence experiments.

### 2.1. Optical Subsystem

Fluorescence excitation is provided by a high-power 470 nm LED (Thorlabs SOLIS-470C) driven by a pulse-modulation LED driver (Thorlabs DC2200). The illumination is shaped using two lenses and a diffuser to maximize illumination uniformity. A bandpass excitation filter (Chroma ET470/40) spectrally filters the illumination to the flavin excitation band.

Fluorescence emission is collected by a Zeiss Milvus 100 mm f/2 M ZF.2 photographic lens. An emission bandpass filter (Chroma ET525/50) is placed between the lens and the camera to suppress excitation light outside of the desired wavelength range. Images are recorded by a back-illuminated scientific sCMOS camera (PCO edge 4.2 bi). The camera is operated in external trigger mode, with frame acquisition initiated by TTL pulses from a National Instruments data acquisition system (DAQ), as described in Section 2 2.4.

The entire optical assembly is mounted on a Thorlabs MB2424 optical breadboard using standard Thorlabs opto-mechanical components (pillar posts, right-angle clamps, SM2 lens tubes, and RM1G cage cubes). The camera and LED source are mounted vertically, pointing downward onto the sample stage, providing a top-illumination, top-collection epifluorescence geometry. This geometry was chosen to eliminate the need for any optical components beneath the sample, leaving the space below the stage accessible for the electromagnet assembly.

Illumination uniformity over the sample footprint was characterized empirically using a calibrated optical power meter at marked positions across a 100 mm Petri dish. Full calibration can be found in Section 3 3.4.

### 2.2. Magnetic Field Subsystem

Three-dimensional magnetic field generation is provided by a GMW Associates Model 5204 vector electromagnet. This commercial electromagnet contains three independently driven coils arranged in a non-coplanar geometry, enabling the creation of an arbitrary field vector at the sample position. The device is water-cooled to permit sustained operation at high current; a Thermo Scientific Thermoflex recirculating chiller provides the required coolant flow. The use of a purpose-built vector electromagnet, rather than a home-wound Helmholtz coil assembly, was driven by the laboratory’s existing access to this instrument; a discussion of more cost-effective alternatives for groups building this instrument from scratch is provided in Section 6 6.2.

Each coil is driven by an analog voltage output from the NI-DAQ system, as described in Section 2 2.4. The voltage-to-field relationship is determined during calibration (Section 3) and is not assumed to be identical across axes or perfectly linear; the 3 × 3 calibration matrix captures cross-axis coupling as well as individual axis sensitivities.

A known limitation of the GMW 5204 geometry is imperfect spatial uniformity over the extended sample volume defined by a 100 mm Petri dish. While the magnetic field at the calibrated center position is controlled with sub-percent accuracy, colonies near the periphery experience different field magnitudes and orientations. This spatial variation is characterized in Section 3 3.4 and its implications for inter-colony comparisons are discussed in Section 6 6.3.

### 2.3. Mechanical Assembly and Sample Stage

The instrument is built on a Thorlabs MB2424 breadboard secured to a mobile cart, providing both a stable optical table surface and the flexibility to relocate the instrument when needed. The electromagnet is mounted on the breadboard using a custom 3D-printed support (big_magnet_support.stl, design files available in the online repository [17]).

A custom sample stage platform (sample_support.stl) is elevated above the electromagnet on four non-magnetic aluminum extrusion legs cut to a length that positions the stage surface at the center of the electromagnet’s field volume without mechanical contact. The stage incorporates two dedicated features: (1) a precisely dimensioned slot for a Lakeshore F71 teslameter probe, which ensures reproducible placement of the magnetometer at the field-center position during calibration and validation, and (2) a registration slot for 100 mm Petri dishes, which ensures that every sample is placed at the same location relative to the field center across experiments. The sample stage is shown in Figure 2, with the axis directions illustrated (following the magnetometer definitions for operational simplicity).

**Fig. 2.**
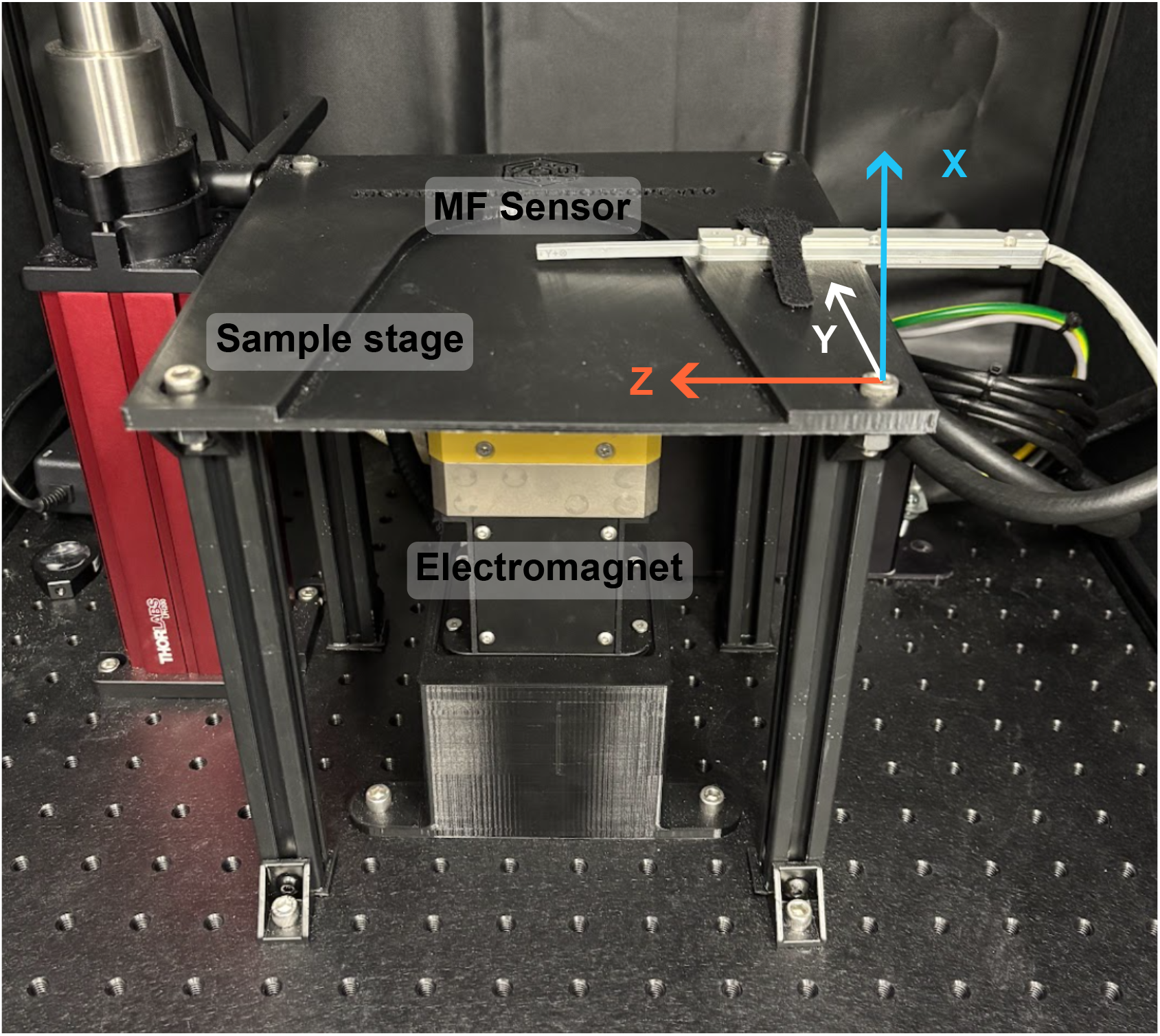
Sample stage. Photograph of the sample stage and magnetic field calibration setup. The biological sample is positioned directly above the electromagnet in a dedicated slot. A three-axis magnetic field (MF) sensor is used to characterize the magnetic field at a specific location. The coordinate system adopted throughout this work is indicated.

To suppress ambient light, the instrument is enclosed in a blackout enclosure constructed from a cage of 2020 aluminum extrusions with rigid black foam panels closing the top and sides. Front and rear access panels use blackout curtain fabric held closed by magnetic and hook-and-loop closures for access convenience. The magnets used for the closures are located sufficiently far from the sample position that their stray field contribution at the sample center is negligible; this was verified by magnetometer measurement during construction.

### 2.4. Electronics and Synchronization

All analog and digital signals are generated and acquired by a National Instruments DAQ system (NI-782258-01) using two data acquisition cards configured in NI MAX (one of these is external, the other one is embedded in the GMW electromagnet electronics). The signal routing is summarized in Table I.

**Table I.**
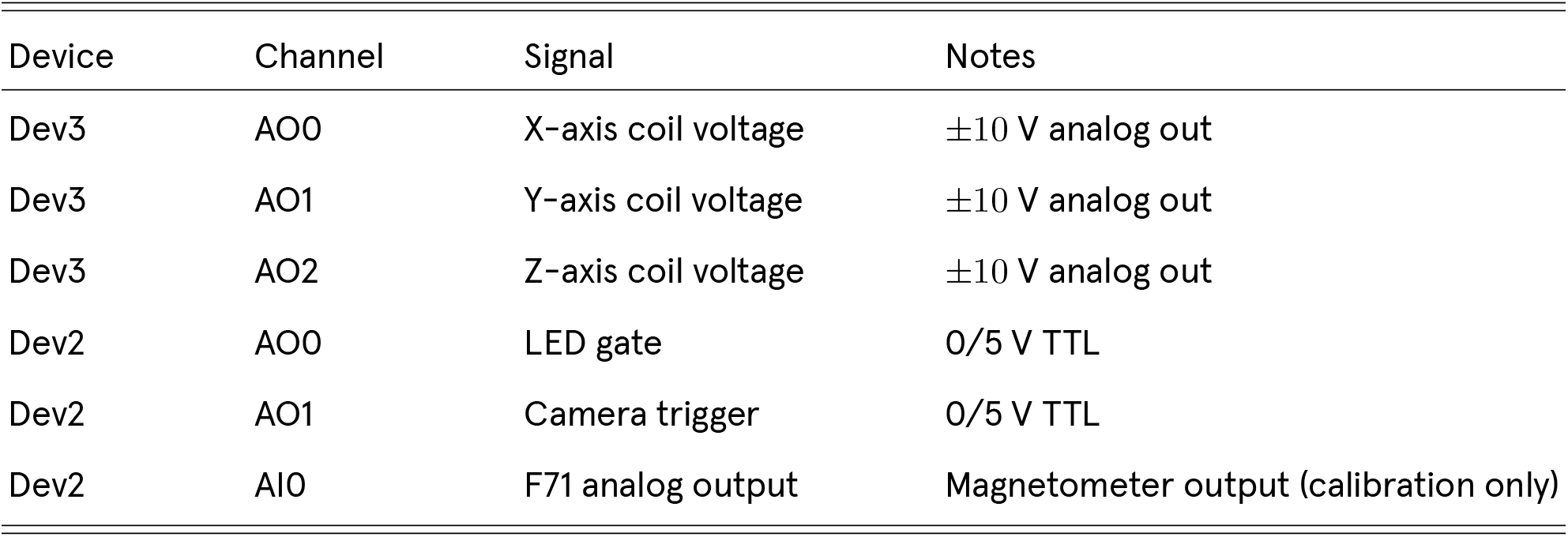
DAQ signal routing.

Waveforms for all output channels are rendered on a common 10 kHz sample clock prior to experiment execution. This shared timebase ensures that the camera trigger, LED gate, and coil voltage waveforms remain precisely aligned throughout the experiment, regardless of its duration. The final sample of each waveform is always forced to zero on all channels, eliminating residual currents in the coils after experiment completion.

The camera is configured in external trigger mode via Micro-Manager [18], controlled from Python using the pycromanager interface [19]. Each camera trigger pulse initiates a single frame acquisition; the LED gate opens a programmable interval before the trigger edge to allow the illumination to stabilize before the exposure begins. The LED advance interval and pulse duration are user-configurable parameters in the experiment GUI (experiment_gui.py) [17].

## 3. Calibration and Field Characterization

Since the scope of this instrument is to investigate the effect of magnetic fields on protein fluorescence, calibration of the magnetic field conditions at the sample location is a crucial part of the design. In this section, different calibration procedures are outlined, covering how the electromagnet is controlled via code, practical software tools that can be used daily as sanity checks, field characterization at the center of the sample location, magnetic field spatial uniformity, and illumination intensity spatial uniformity.

### 3.1. Coil Calibration Procedure

The relationship between coil drive voltages and the resulting magnetic flux density at the sample position is described by a linear model:

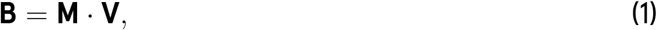

where **B** ∈ R^3^ is the flux density vector (in mT) at the sample center, **V** ∈ R^3^ is the vector of coil drive voltages (in V), and **M** ∈ R^3^*^×^*^3^ is the calibration matrix. The matrix **M** captures both the individual coil sensitivities (diagonal entries) and any cross-axis coupling (off-diagonal entries). The inverse relationship, **V** = **M***^−^*^1^ · **B**, is used at runtime to convert a user-specified target field vector into coil drive voltages.

Calibration is performed using the calibration_gui.py software module, with a Lake Shore F71 teslameter probe placed in its dedicated slot on the sample stage. The calibration routine applies a set of voltages while reading the teslameter response, and fits the resulting measurements by least squares to determine **M**. The ambient field **B**_0_ — the flux density measured at **V** = 0 — is also recorded and saved separately; it can optionally be subtracted from target field requests at runtime to compensate for the local geomagnetic field and any remanent magnetization in the electromagnet. The result of the calibration procedure, along with the predicted behavior calculated using **M**, is reported in Figure 3a.

**Fig. 3.**
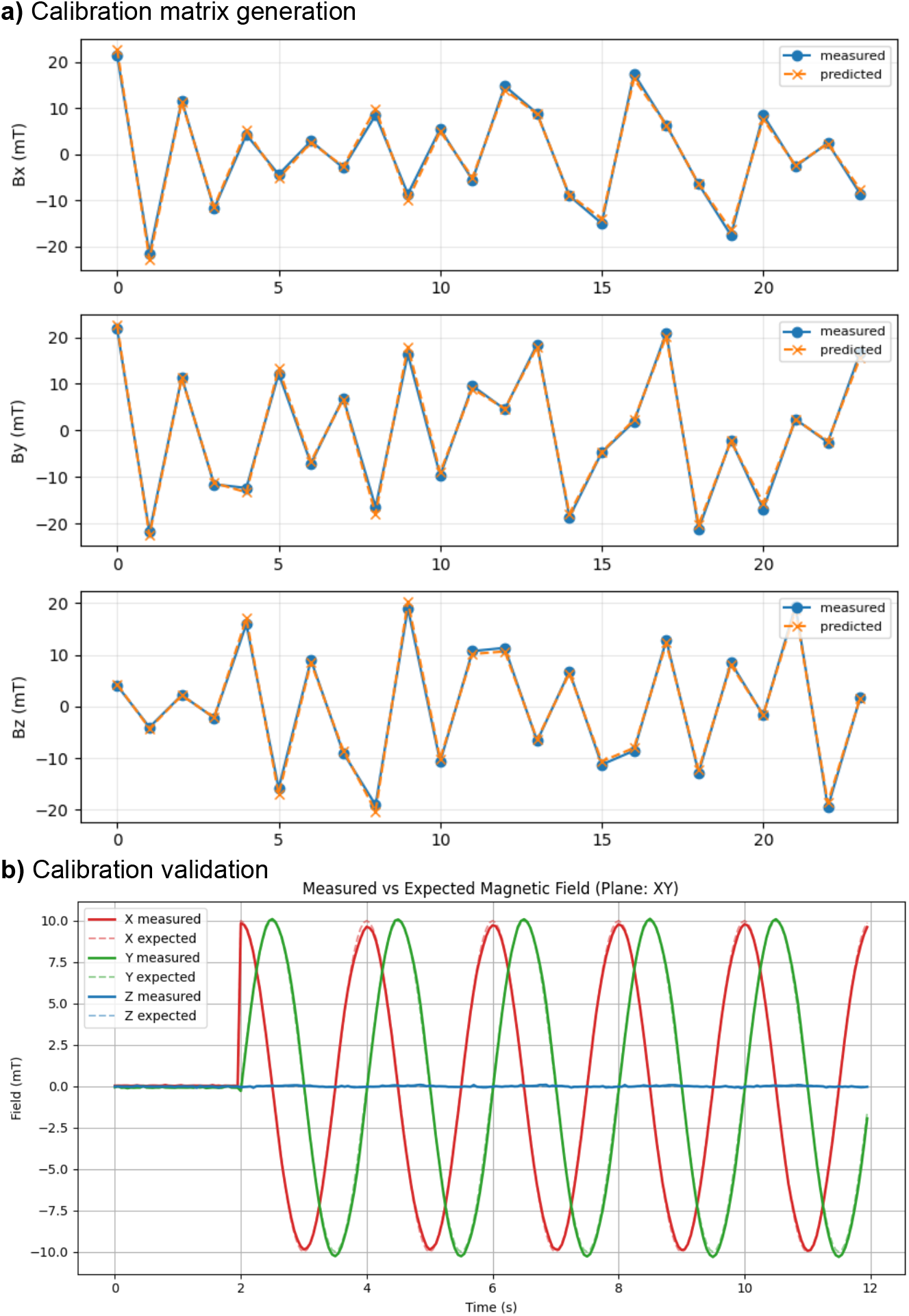
Calibration and validation of programmable vector magnetic field generation. (a) Calibration matrix generation using measurements of 24 arbitrary magnetic field vectors spanning the accessible operating range of the electromagnet. Predicted magnetic field components obtained from the fitted calibration matrix are compared with independent measurements for each Cartesian axis, demonstrating accurate reconstruction of the applied fields. (b) Validation of the calibrated system through generation of a 10 mT rotating magnetic field in the XY plane. Solid lines indicate the magnetic field measured with the three-axis magnetometer, while dashed lines show the expected field trajectory calculated from the programmed waveform. The close agreement between measured and expected fields confirms accurate dynamic vector field generation for magnetofluorescence experiments.

### 3.2. Static Field Accuracy

Static field accuracy was validated using field_test_gui.py, which applies the inverse calibration to a set of target field vectors and compares the resulting measured field (as read by the teslameter) to the target. This GUI was designed to provide an easy and quick daily sanity check, to ensure magnetic field control is behaving as expected and stable over time.

### 3.3. Dynamic Field Characterization

Dynamic field performance was evaluated using field_behavior_gui.py, which imposes rotating and ramp field sequences while recording the teslameter output. To illustrate the performance of the calibration, an example is reported in Figure 3b, where a rotating field in the XY plane of amplitude 10 mT was applied at a frequency of 0.5 Hz. The measured magnetic field behavior (solid lines) closely follows the expected behavior (dashed lines), thus demonstrating good control of the magnetic field in terms of timing and amplitude. The dynamic range has not been extensively tested, since the experimental protocols consist of ON/OFF cycles that hold a constant field over several seconds (see examples in Section 5), making any extensive dynamic calibration beyond the scope of this work.

### 3.4. Spatial Field Uniformity

The GMW Associates 5204 electromagnet is designed for single-point field generation and does not guarantee a uniform field over an extended sample volume. To quantify the spatial variation of |*B*| across the 100 mm Petri dish footprint, a Lake Shore F71 teslameter probe was manually positioned at a grid of points across the sample stage, corresponding to the locations marked as white dots in Figure 4, and measurements of *B_x_*, *B_y_*, *B_z_*, and total field magnitude |*B*| were recorded at each. A fixed coil drive corresponding to a target field of 10 mT at the center along the out-of-plane (X) direction was applied; qualitatively similar spatial profiles were observed at other field magnitudes. The measured field values were fitted using a two-dimensional quadratic surface, which was subsequently used to estimate the spatial field variation across the imaging area. Within the central 40 mm diameter region used for quantitative analysis, the fitted magnetic field magnitude was 95.5 ± 4.6% of the central value.

**Fig. 4.**
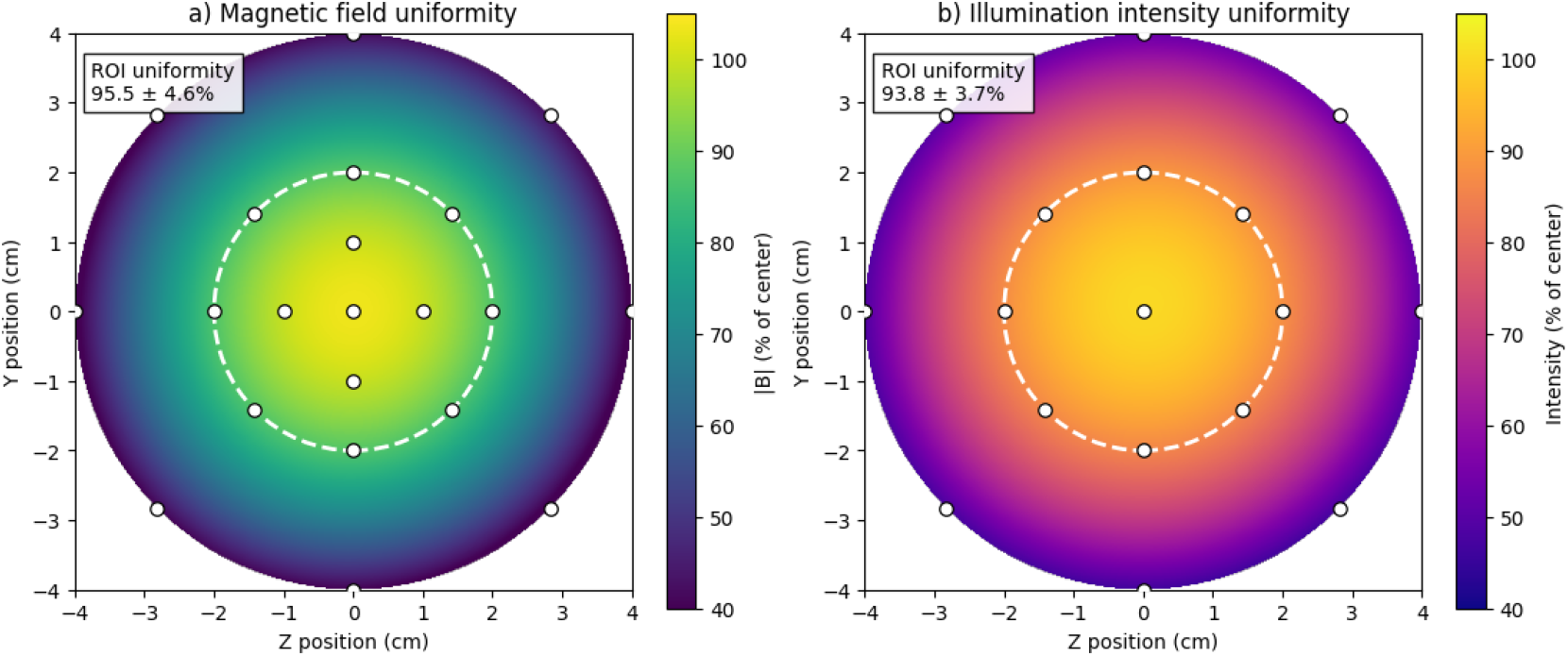
Spatial characterization of magnetic field and illumination uniformity across the sample plane. (a) Magnetic field magnitude, normalized to the field at the center of the sample stage, measured using a three-axis magnetometer and interpolated using a two-dimensional quadratic surface fit. (b) Excitation illumination intensity, normalized to the intensity at the center of the field of view, measured using a Thorlabs S120C photodiode sensor and interpolated using the same fitting procedure. White circles indicate the measurement locations used for calibration, while the dashed circle denotes the 4 cm diameter region of interest (ROI) used for quantitative analysis. Within this ROI, the magnetic field magnitude and illumination intensity remained 95.5 ± 4.6% and 93.8 ± 3.7% of their respective central values (mean ± standard deviation of the fitted surface).

**Fig. 5.**
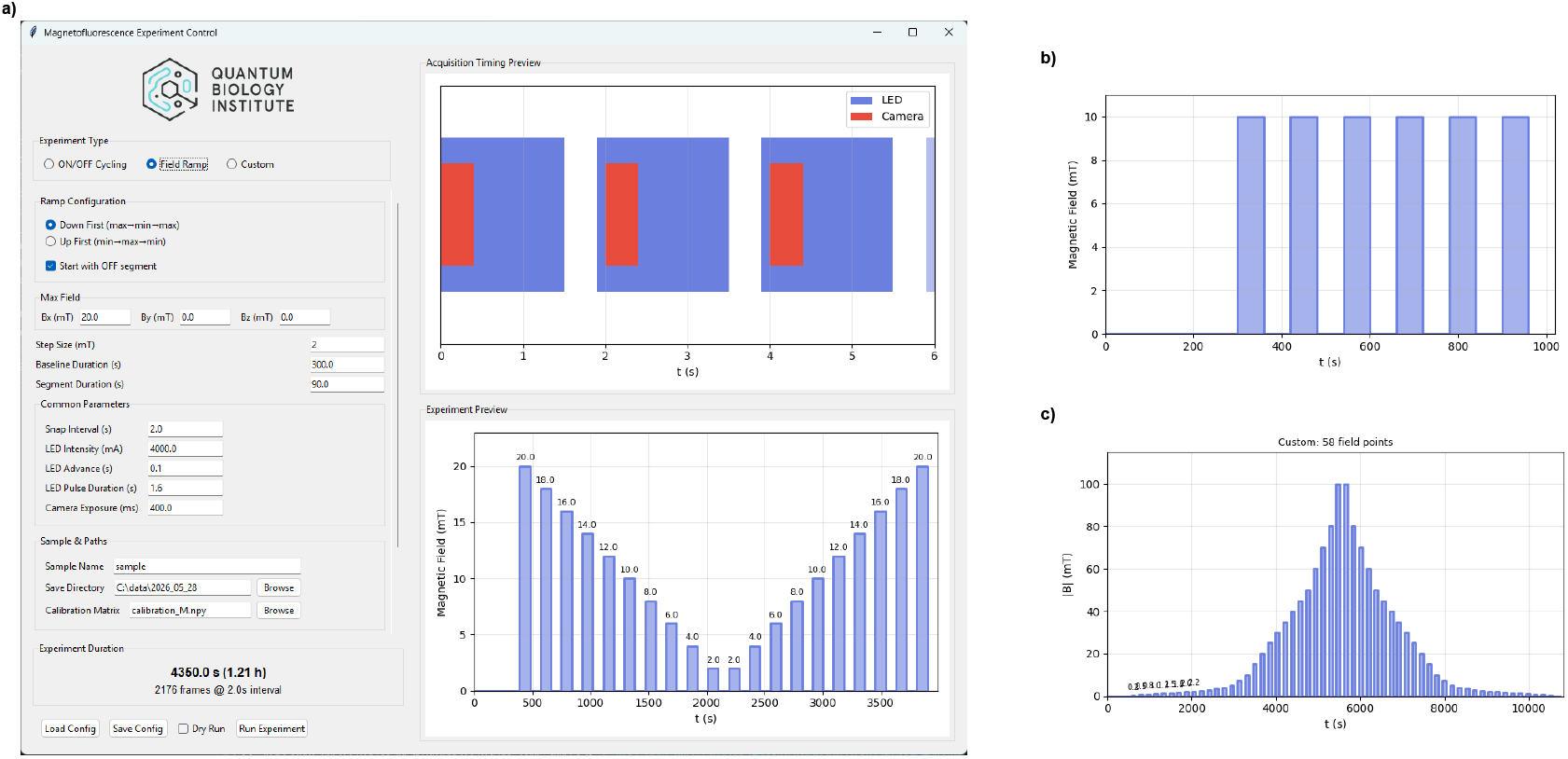
Graphical user interface for experiment configuration and representative magnetic field protocols. (a) Main control interface showing configuration of magnetic field parameters, synchronized LED illumination and camera acquisition, and real-time previews of acquisition timing and the programmed magnetic field schedule. The interface shown is configured for a field ramp experiment. (b) Example ON/OFF cycling protocol used for time-resolved magnetofluorescence measurements. (c) Example user-defined magnetic field schedule illustrating arbitrary programming of magnetic field magnitude as a function of time.

Excitation illumination uniformity was characterized at the same marked locations using a Thorlabs analog handheld laser power meter console equipped with an S120C photodiode sensor, with the LED source operated at full power. The S120C sensor has an active aperture diameter of 9.5 mm, corresponding to an effective collection area of 0.709 cm^2^; measured optical powers were converted to excitation intensities by dividing by the sensor area. At full LED output, the excitation intensity at the center of the field of view was 6.63 mW/cm^2^, corresponding to the 100% reference level shown in Figure 4b. As with the magnetic field characterization, the measured illumination profile was fitted using a two-dimensional quadratic surface. Within the central 40 mm diameter analysis region, the fitted excitation intensity was 93.8 ± 3.7% of the central value.

To minimize systematic variation arising from both magnetic field and illumination nonuniformity, quantitative MFE analysis was restricted to colonies located within this central 40 mm diameter region.

In summary, the magnetic field at the sample stage can be controlled in an automated way via Python code using a calibration matrix. The accuracy and uniformity have been characterized extensively through a systematic procedure. The light intensity uniformity over the sample area has also been carefully characterized.

## 4. Control Software

The control software package was designed to provide a unified interface for instrument calibration, magnetic field generation, experiment execution, and data acquisition. The software architecture consists of a collection of specialized graphical user interfaces (GUIs) that share a common hardware abstraction layer and experiment engine. This modular architecture simplifies maintenance, promotes code reuse, and ensures that calibration parameters are applied consistently across all experimental workflows. Together, these software components enable users with minimal experience in magnetic instrumentation to perform fully automated magnetofluorescence experiments through an intuitive graphical interface, while retaining the flexibility required for advanced custom protocols.

### 4.1. Architecture

The control software is written in Python and organized around two shared modules that implement all hardware interaction, with a set of Tkinter graphical user interfaces layered on top for day-to-day operation. The two core modules are:

- experiment_runners.py — the experiment engine, responsible for waveform generation, DAQ thread management, Micro-Manager acquisition control, and per-frame metadata writing.
- hardware_io.py — hardware wrappers providing a CoilDriver class for NI-DAQ analog output, F71Analog and F71SCPI classes for teslameter readout, and calibration-loading utilities.

This two-module design ensures that the hardware abstraction layer is shared across all GUI applications, eliminating code duplication and ensuring that calibration is applied consistently regardless of which GUI is in use.

### 4.2. Waveform Generation and Experiment Modes

Three experiment modes are implemented, each with a dedicated waveform builder and orchestrator function:

#### ON/OFF Cycling

A square-wave protocol with a configurable baseline period followed by *N* alternating ON and OFF segments, each of fixed duration, at a user-specified target field vector. This mode is intended for standard ON/OFF measurements.

#### Field Ramp

A staircase protocol that steps the field magnitude from a maximum to a minimum (or vice versa) across *N* levels along a specified direction vector, with optional OFF segments interleaved between steps. This mode is designed for dose-response characterization of field-magnitude dependence.

#### Custom Sequence

An arbitrary field sequence loaded from a comma-separated value (CSV) file specifying (*B_x_, B_y_, B_z_*) triplets in mT, one per row. This mode enables arbitrary user-defined field sequences.

All three waveform builders render their output on a common 10 kHz time grid, producing three synchronized output arrays: coil voltages for the three DAQ analog output channels, the LED gate TTL waveform, and the camera trigger TTL waveform. The LED gate is opened at a user-configurable lead time before each camera trigger edge, allowing the LED driver to reach steady state before the exposure begins.

When a requested field vector requires coil drive voltages outside the ±10 V DAQ output range, the vector is uniformly scaled down to the largest feasible magnitude and the corresponding frames are flagged as saturated: true in the per-frame metadata. This behavior prevents silent field errors while preserving the direction and relative magnitude of the requested vector.

### 4.3. Data Acquisition and Metadata

Experiment execution is managed by two concurrently running threads: a DAQ thread that plays the pre-rendered finite waveforms via NI-DAQmx, and an acquisition thread that drives Micro-Manager to capture frames. The threads are synchronized by the shared hardware timebase; the acquisition thread uses the camera’s external trigger mode so that each frame begins exactly at the corresponding trigger edge in the DAQ waveform.

Per-frame metadata is generated during waveform construction and injected into each OME-TIFF frame’s userData field by the acquisition thread. Each frame record includes frame_index, planned_time_s, field_on (boolean), magneticField_mT (3-vector), coilVoltages_V (3-vector), and saturated (boolean flag). The same metadata is also written to an experiment_summary.json file in the output directory. Embedding metadata in the image file ensures that the experimental conditions are permanently coupled to the image data and cannot be accidentally separated during archiving or sharing.

### 4.4. Software Availability

The complete software package, including all GUI applications, shared modules, example field sequence files, and conda and pip environment specifications, is freely available at https://github.com/Quantum-Biology-Institute/Bacterioscope [17] under the Apache License 2.0. Detailed build documentation and assembly instructions are provided in publicly accessible electronic laboratory notebooks at https://data.quantumbiology.org under Research: Biology LA / Bacterioscope / Chapter 1: ELN.

## 5. Demonstration: Detection of Magnetic Field Effects in Live Bacterial Cells

To demonstrate functionality, the instrument was used to image *E. coli* colonies expressing the engineered magnetosensitive fluorescent protein MagLOV2 [13–15] under programmable magnetic field stimulation protocols. Full biological analysis and interpretation are reported separately in Ref. [15]; here, we focus on representative experiments illustrating the capabilities of the imaging platform.

Fluorescence images of bacterial plates containing hundreds of spatially separated colonies were acquired under both ON/OFF cycling and magnetic field ramp protocols. A representative raw fluorescence image is reported in Figure 6a. In ON/OFF experiments, alternating magnetic field segments were applied while continuously imaging fluorescence emission, enabling direct observation of field-dependent fluorescence changes in individual colonies. Representative traces are shown in Figure 6, where reproducible fluorescence modulation is observed upon application and removal of the external magnetic field. In this case, it was demonstrated that, for the same bacterial colony, the MFE magnitude can be reversed for different magnetic fields, showing a non-monotonic dependence between fluorescence intensity change and magnetic field amplitude.

**Fig. 6.**
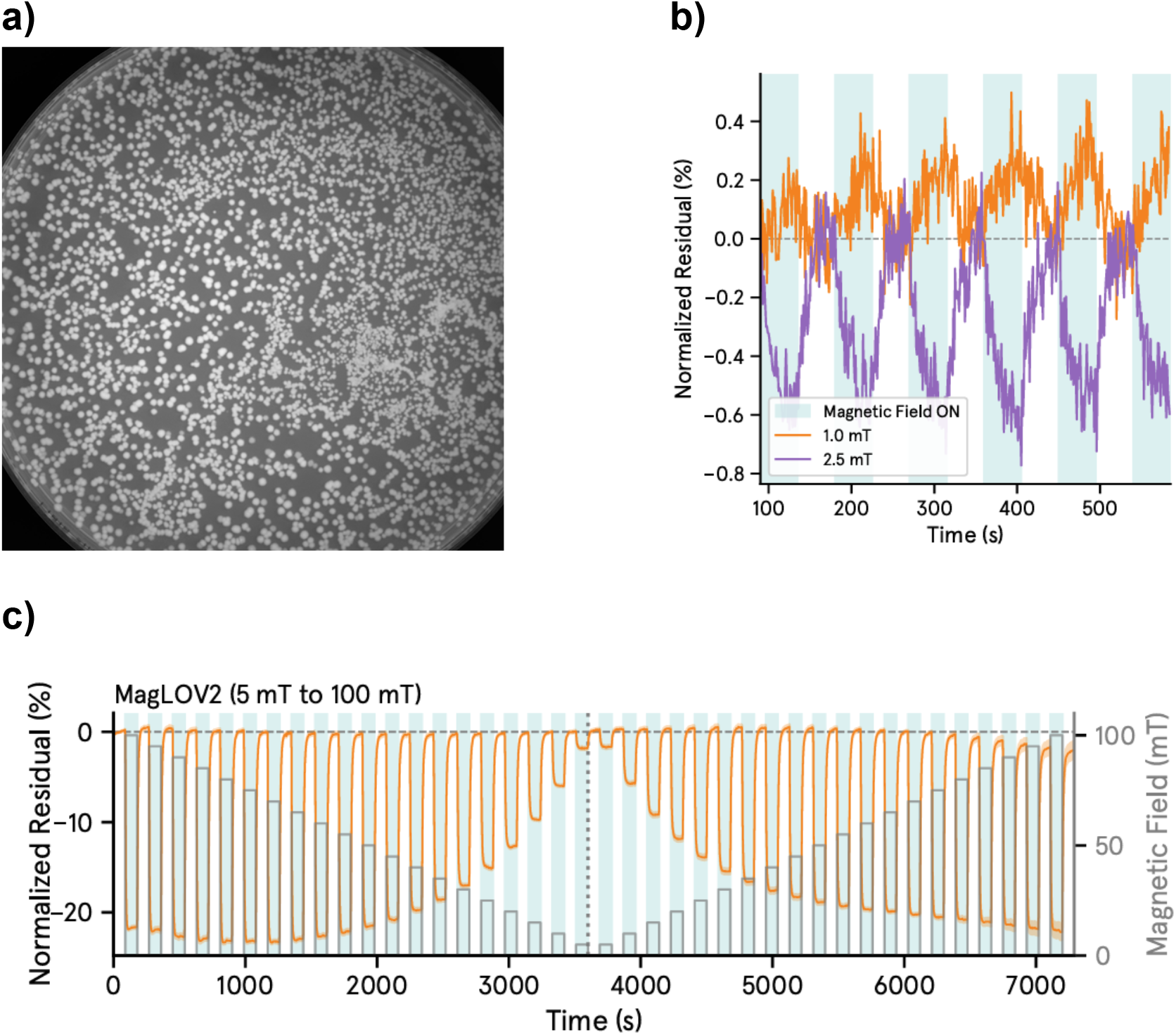
Demonstration of instrument performance using the magnetosensitive protein MagLOV2, expressed in E. coli. (a) Representative fluorescence image acquired during an experiment. (b) Example ON/OFF magnetofluorescence experiment comparing fluorescence responses to 1.0 and 2.5 mT magnetic fields in the same bacterial colony. Shaded regions indicate periods during which the magnetic field was applied. (c) Representative field-ramp experiment in which the magnetic field magnitude was varied between 5 and 100 mT using repeated ON/OFF cycles. The measured fluorescence response (orange) is shown together with the applied magnetic field protocol (gray bars), illustrating the dependence of the magnetic field effect on field strength. These experiments demonstrate the ability of this instrument to perform automated time-resolved magnetofluorescence measurements.

To demonstrate programmable waveform control, the instrument was additionally operated in ramp mode, in which the applied magnetic field magnitude was stepped across a user-defined sequence while fluorescence images were continuously acquired. Representative ramp experiments are shown in Figure 6c, illustrating stable long-duration synchronized acquisition during dynamic magnetic field exposure.

Fluorescence images were automatically segmented into individual colonies using rolling-ball background subtraction [20] and Otsu thresholding [21] with watershed separation [22]. The code to perform this analysis is available at https://github.com/ Quantum-Biology-Institute/mfe_fit_analysis [23]. Mean fluorescence intensity was extracted for each colony following local background subtraction. Magnetic field effects were quantified by fitting fluorescence traces during field-OFF segments with a multi-exponential baseline model and computing normalized residuals relative to the fitted baseline. These residuals provide a measurement of the magnetic field effect.

To demonstrate the applicability of this instrument for investigating proteins in solution and biological assays, cell lysates from *E. coli* expressing MagLOV2 were imaged during repeated cycles of 100 mT magnetic field ON/OFF cycling (Figure 7a). The corresponding normalized residual mean fluorescence intensity of the wells is shown in Figure 7b.

**Fig. 7.**
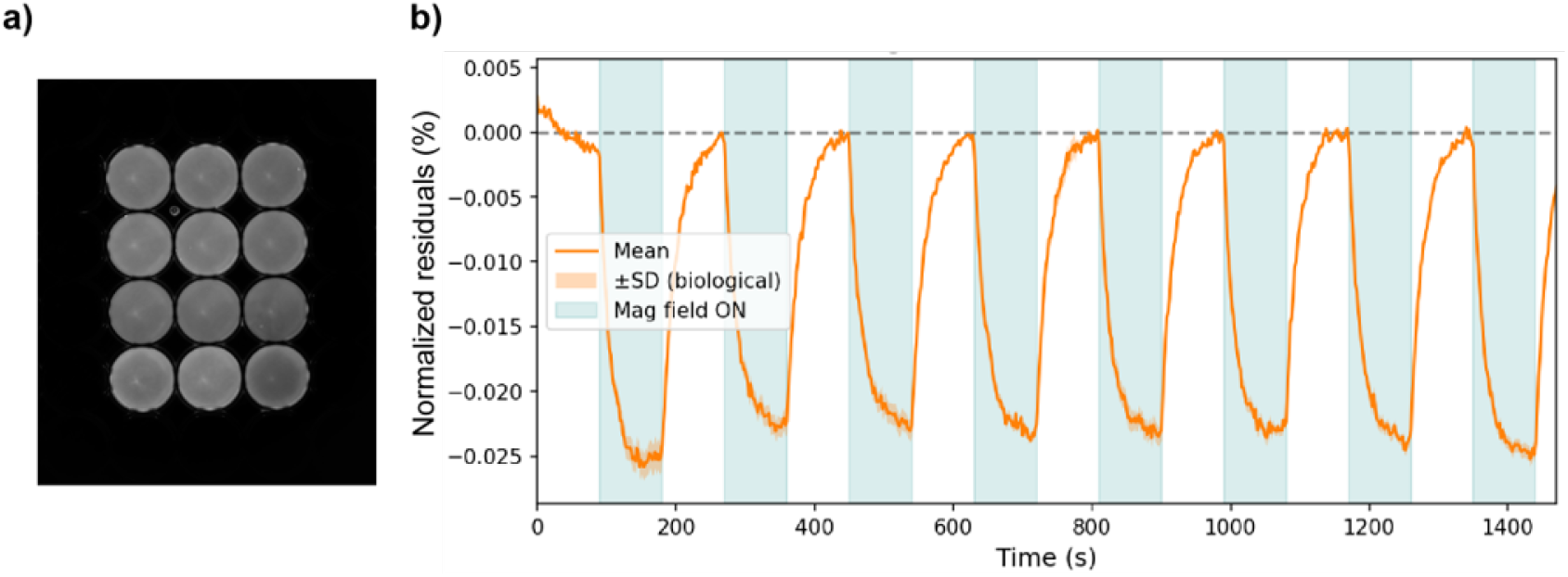
Demonstration of magnetofluorescence measurements in protein solutions. (a) Representative fluorescence image of cell lysates from *E. coli* expressing MagLOV2 loaded into a 96-well plate lid. (b) Normalized residual fluorescence traces extracted from the wells during repeated 100 mT magnetic field ON/OFF cycles. These measurements demonstrate that the instrument is readily applicable not only to bacterial colony imaging but also to fluorescence assays performed in multi-well formats, extending its utility to purified proteins, cell lysates, and biochemical reaction mixtures.

These measurements demonstrate that this setup can reliably detect magnetic field effects as low as ∼0.1% across hundreds of bacterial colonies and fluorescent proteins in solutions or reactions simultaneously, while supporting flexible programmable magnetic stimulation protocols.

## 6. Discussion

### 6.1. Summary of Instrument Capabilities

Table II summarizes the key performance specifications of this screening setup.

**Table II.** Instrument key specifications.

| Parameter | Value |
| --- | --- |
| Camera | PCO edge 4.2 bi (sCMOS) |
| Excitation wavelength | 470/40 nm (Chroma ET470/40) |
| Emission band | 525/50 nm (Chroma ET525/50) |
| Field of view | 80 mm × 80 mm |
| Magnetic field range (per axis) | ±100 mT |
| Static field accuracy | < 1% RMS |
| Field uniformity across 40 mm diameter ROI | 95.5 ± 4.6% |
| Illumination uniformity across 40 mm diameter ROI | 93.8 ± 3.7% |
| Synchronization timebase | 10 kHz |
| Approximate build cost | 70,000 USD<sup>a</sup> |

<sup>a</sup> Cost can be substantially reduced by replacing the electromagnet with a home-built Helmholtz coil system.
Other cost-cutting ideas are discussed in Section 6.6.4.

### 6.2. Comparison to Existing Approaches

This instrument was inspired by Andrew G. York’s chicken blaster setup [24]. His imaging system was later repurposed to perform magnetofluorescence experiments, where a magnetic field was applied by sliding permanent magnets under the sample. Such an instrument was used to screen for fluorescence sensitivity to external magnetic fields, resulting in the MagLOV2 protein [13]. This instrument shares much of its hardware with that setup but implements automated experiments with programmable control over magnetic field magnitude and direction across a broad operating range.

Other setups used in the scientific literature for investigating MFEs in biology include microscopy setups, such as confocal microscopy [25]. This instrument trades spatial resolution for throughput: by imaging at colony rather than cellular resolution, it can monitor hundreds of genetically distinct colonies simultaneously under similar field conditions, making it well suited for screening applications in synthetic biology and quantum biology research.

### 6.3. Limitations and Mitigations

The primary limitation of the current instrument is the spatial non-uniformity of the GMW 5204 electromagnet over the 100 mm dish footprint (Section 3 3.4). Colonies at the periphery of the dish experience a different field magnitude than those at the center, which introduces a systematic positional confound into cross-colony comparisons. This can be partially mitigated by restricting quantitative comparisons to colonies within a 40 mm radius of the dish center, or by applying a spatial correction using the measured uniformity map. Future builds may benefit from a purpose-wound, water-cooled Helmholtz coil assembly that might provide substantially more uniform fields over the required volume, at the expense of a lower field intensity ceiling and fixed directionality. This would also reduce the overall cost and complexity of the instrument.

A second limitation is that the current optical configuration provides uneven illumination over a large area of a 100 mm dish. This can also be overcome by restricting quantitative analysis to a 40 mm radius around the center, and/or replacing some of the optical components to achieve better illumination uniformity.

### 6.4. Cost and Accessibility

The dominant cost driver in the current build is the GMW Associates 5204 electromagnet, which was available in the laboratory from prior work. For groups building from scratch, a custom water-cooled Helmholtz coil assembly using off-the-shelf magnet wire and a standard laboratory power supply interfaced via a DAQ analog output can achieve fields on the order of tens of millitesla at a fraction of the cost, with the additional benefit of improved spatial uniformity. While the NI-DAQ hardware represents a secondary cost, lower-cost DAQ options exist that satisfy the minimum channel requirements (three analog outputs for the coils, two TTL outputs for the LED and camera trigger, one analog input for the magnetometer) at reduced cost, at the expense of somewhat more complex wiring. The optical and camera components constitute the remaining major expense; the PCO edge 4.2 bi is a premium scientific camera, and groups for whom the primary cost driver is the camera may find that a less expensive sCMOS camera with comparable quantum efficiency provides adequate performance for colony-level fluorescence measurements.

The approximate total cost of the instrument as built is approximatey 70,000 USD in 2026. A detailed bill of materials is provided in Appendix A.

## 7. Conclusion

We have described an open-source platform for plate-scale magnetofluorescence imaging in live bacterial colonies. The instrument integrates programmable three-axis magnetic field generation with synchronized fluorescence imaging and automated acquisition software, enabling high-throughput screening of magnetic field effects across hundreds of colonies simultaneously.

The instrument’s utility was demonstrated by detecting and quantifying magnetic field effects on the magnetosensitive fluorescent protein MagLOV2 in live colonies of *E. coli*, confirming its value for quantum-biology screening at biologically relevant scales. All design files, build documentation, and control software are made openly available to support replication and adaptation by the scientific community.

This instrument provides a standardized and quantitatively characterized platform for magnetofluorescence measurements, lowering the technical barrier to magnetic field experimentation while promoting reproducibility and cross-laboratory comparability. We anticipate that instruments of this kind will facilitate the systematic investigation of magnetic field effects and contribute to the establishment of experimental standards needed for the continued advancement of magnetobiology.

## Code availability

The control and acquisition software described here is openly available at https://github.com/<OWNER>/Bacterioscope_code_share under the Apache License 2.0.[17] Calibration data files are released under CC0-1.0.

## Appendix A. Bill of materials

**Table A1.** Bill of materials. Approximate component costs (USD). The teslameter is listed for calibration only and is not in the total.

| Component | Model / part number | Supplier | Approx. cost (USD) |
| --- | --- | --- | --- |
| Camera | PCO edge 4.2 bi | Excelitas/PCO | 14,100.00 |
| Objective lens | Zeiss Milvus 100 mm f/2 ZF.2 | B&H / Zeiss | 1,918.00 |
| Excitation filter | Chroma ET470/40 | Chroma | 365.00 |
| Emission filter | Chroma ET525/50 | Chroma | 365.00 |
| LED source | Thorlabs SOLIS-470C | Thorlabs | 1,556.77 |
| LED driver | Thorlabs DC2200 | Thorlabs | 2,593.01 |
| Illumination lens | Thorlabs AC508-100-A-ML | Thorlabs | 174.14 |
| NI-DAQ | NI 782258-01 | National Instruments | 5,351.62 |
| Vector electromagnet | GMW Associates 5204 | GMW Associates | 33,290.00 |
| Recirculating chiller | Thermo Scientific Thermoflex | Fisher Scientific | 8,980.00 |
| Teslameter | Lakeshore F71 | Lakeshore | Calibration only |
| Optical breadboard | Thorlabs MB2424 | Thorlabs | 602.27 |
| Opto-mech. components | Thorlabs posts, clamps, cubes, SM2 tubes (various) | Thorlabs | 1,000.00 |
| Enclosure frame | 2020 aluminum extrusion + corner brackets | Amazon / various | 100.00 |
| Enclosure panels | Black foam core board 24" × 36" | Amazon | 69.99 |
| 3D-printed parts | big_magnet_support.stl, sample_support.stl | In-house / service | 10.00 |
| **Total** |  |  | **70,475.80** |

## 8. Author Contributions

Following the CRediT taxonomy [26]:

1. Alessandro Lodesani: conceptualization; data curation; formal analysis; investigation; methodology; software; validation; visualization; and writing (original draft)
2. Brian L. Ross: investigation; validation; and writing (review and editing)
3. Venkatesh Sridharan: investigation; validation; and writing (review and editing)
4. Clarice D. Aiello: conceptualization; funding acquisition; validation; and writing (review and editing)

## Acknowledgements

We thank Andrew York for providing a blueprint for a bacterial plate imaging system that inspired this instrument and for helpful discussion around it. We also thank Maria Ingaramo and Nonfiction Laboratories for providing the pRSET-MagLOV2 construct as well as valuable insights into its expression and imaging. We would like to thank Morgan Sosa for her assistance with formatting, revising, and editing the manuscript, and we thank Michael Montague, Todd Thaxton, and Bret Gaitan for critically reading the manuscript and providing helpful feedback. We also acknowledge the use of Claude Code for developing the python code and graphical user interfaces.

## Footnotes

The [Quantum Biology Institute](http://quantumbiology.org/) is a California non-profit 501(c)(3) focused research organization that performs basic research underpinning the quantum biology field in an open-science fashion.

---

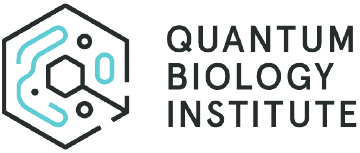

## References

[1] P. J. Hore and H. Mouritsen, The radical-pair mechanism of magnetoreception, Annual Review of Biophysics 45, 299 (2016).

[2] H. Zadeh-Haghighi and C. Simon, Magnetic field effects in biology from the perspective of the radical pair mechanism, Journal of the Royal Society Interface 19, 20220325 (2022).

[3] A. Lodesani, G. Anders, L. Bougas, T. Lins, D. Budker, P. Fierlinger, and C. D. Aiello, Weak magnetic field effects in biology are measurable—accelerated *Xenopus* embryogenesis in the absence of the geomagnetic field, bioRxiv 10.1101/2024.10.10.617626 (2024).

[4] M. Montague, A. Lodesani, and C. D. Aiello, Escherichia coli K12 exhibits a ∼50% longer lag phase, but no difference in log phase growth rate, under hypomagnetic conditions (19 nT) 10.64898/2026.04.13.717819 (2026).

[5] E. W. Evans, C. A. Dodson, K. Maeda, T. Biskup, C. J. Wedge, and C. R. Timmel, Magnetic field effects in flavoproteins and related systems, Interface Focus 3, 20130037 (2013).

[6] K. B. Henbest et al., Magnetic-field effect on the photoactivation reaction of Escherichia coli DNA photolyase, Proceedings of the National Academy of Sciences 105, 14395 (2008).

[7] U. E. Steiner and T. Ulrich, Magnetic field effects in chemical kinetics and related phenomena, Chemical Reviews 89, 51 (1989).

[8] C. D. Aiello, B. L. Ross, A. Lodesani, and M. L. Sosa, A physicist-friendly primer on the hamiltonian for quantum sensing in proteins: analytical expressions and insights for a toy model of the radical-pair mechanism, (2026), arXiv:2604.18608 [physics.bio-ph].

[9] D. R. Kattnig et al., Chemical amplification of magnetic field effects relevant to avian magnetoreception, Nature Chemistry 8, 384 (2016).

[10] J. Xu et al., Magnetic sensitivity of cryptochrome 4 from a migratory songbird, Nature 594, 535 (2021).

[11] K. Maeda et al., Magnetically sensitive light-induced reactions in cryptochrome are consistent with its proposed role as a magnetoreceptor, Proceedings of the National Academy of Sciences 109, 4774 (2012).

[12] C. K. Austvold, S. M. Keable, M. Procopio, and R. J. Usselman, Quantitative measurements of reactive oxygen species partitioning in electron transfer flavoenzyme magnetic field sensing, Frontiers in Physiology 15, 1348395 (2024).

[13] R. F. Hayward, A. Rai, J. R. Lazzari-Dean, A. E. Y. T. Lefebvre, A. G. York, and M. Ingaramo, Magnetic control of the brightness of fluorescent proteins (2024).

[14] G. Abrahams, A. Štuhec, V. Spreng, R. Henry, I. Kempf, J. James, K. Sechkar, S. Stacey, V. Trelles-Fernandez, L. M. Antill, C. R. Timmel, J. J. Miller, M. Ingaramo, A. G. York, J.-P. Tetienne, and H. Steel, Quantum spin resonance in engineered proteins for multimodal sensing, Nature 649, 1172 (2026).

[15] B. L. Ross, A. Lodesani, and C. D. Aiello, The magnetic field-dependent fluorescence of MagLOV2 in live bacterial cells is consistent with the radical pair mechanism 10.64898/2026.02.18.706690 (2026).

[16] K. M. Xiang, H. Lampson, R. F. Hayward, A. G. York, M. Ingaramo, and A. E. Cohen, Mechanism of giant magnetic field effect in a red fluorescent protein, Journal of the American Chemical Society 147, 18088 (2025).

[17] A. Lodesani and M. Sosa, Bacterioscope: Control and acquisition code, https://github.com/<OWNER>/Bacterioscope_code_share (2026), accessed: 2026-07-24.

[18] A. D. Edelstein et al., Advanced methods of microscope control using Micro-Manager software, Journal of Biological Methods 1, e10 (2014).

[19] H. Pinkard et al., Pycro-manager: open-source software for customized and reproducible microscope control, Nature Methods 18, 226 (2021).

[20] S. R. Sternberg, Biomedical image processing, Computer 16, 22 (1983).

[21] N. Otsu, A threshold selection method from gray-level histograms, IEEE Transactions on Systems, Man, and Cybernetics 9, 62 (1979).

[22] L. Vincent and P. Soille, Watersheds in digital spaces: An efficient algorithm based on immersion simulations, IEEE Transactions on Pattern Analysis and Machine Intelligence 13, 583 (1991).

[23] B. L. Ross, mfe_fit_analysis: Colony fluorescence segmentation and magnetic field effect analysis, https://github.com/Quantum-Biology-Institute/mfe_fit_analysis (2026), accessed: 2026-07-28.

24. [24] https://andrewgyork.github.io/relaxation_sensors/appendix.html#chicken_blaster.

[25] V. Déjean, M. Konowalczyk, J. Gravell, M. J. Golesworthy, C. Gunn, N. Pompe, O. Foster Vander Elst, K.-J. Tan, M. Oxborrow, D. G. A. L. Aarts, S. R. Mackenzie, and C. R. Timmel, Detection of magnetic field effects by confocal microscopy, Chemical Science 11, 7772 (2020).

[26] CRediT – Contributor Roles Taxonomy, https://credit.niso.org/, accessed: 2026.

